# The Endoplasmic Reticulum Membrane Complex Promotes Proteostasis of GABA_A_ Receptors

**DOI:** 10.1101/2022.03.03.482920

**Authors:** Angela Whittsette, Ya-Juan Wang, Ting-Wei Mu

**Affiliations:** Department of Physiology and Biophysics, Case Western Reserve University School of Medicine, 10900 Euclid Avenue, Cleveland, Ohio 44106, USA

## Abstract

The endoplasmic reticulum membrane complex (EMC) plays a critical role in the biogenesis of tail-anchored and a subset of multi-pass membrane proteins in the endoplasmic reticulum. However, due to the nearly exclusive expression of neurotransmitter-gated ion channels in the central nervous system, the role of the EMC in their biogenesis is not well understood. In this study, we demonstrated that the EMC positively regulates the surface trafficking and thus function of endogenous γ-aminobutyric acid (GABA_A_) receptors, the primary inhibitory ion channels in the mammalian brain. Further, among ten EMC subunits, EMC3 and EMC6 have the most prominent effects, indicating a subunit-specific contribution. EMC3 and EMC6 show endogenous interactions with major neuroreceptors, which depends on their transmembrane domains. Overexpression of EMC3 and EMC6 is sufficient to restore the function of epilepsy-associated GABA_A_ receptor variants, suggesting that operating EMC has the potential to ameliorate neurological diseases associated with protein conformational defects.

**In brief:** The multi-subunit EMC serves as an insertase for a subset of membrane proteins and enables their biogenesis in the endoplasmic reticulum. However, the subunit-specific effect of the EMC on multi-pass neuroreceptors is not well understood. Whittsette et al. demonstrate that EMC3 and EMC6 interact with GABA_A_ receptors and positively regulate their trafficking and function.

**Highlights:**

- EMC3 and EMC6 positively regulate the function of endogenous GABA_A_ receptors.
- The EMC interacts with major endogenous neuroreceptors.
- The interaction between EMC and GABA_A_ receptors depends on the EMC transmembrane domains.
- Overexpressing the EMC is sufficient to restore the function of pathogenic GABA_A_ receptor variants.

## INTRODUCTION

Acquiring correct transmembrane topology is essential for the function of membrane proteins, which consist of about 30% of the eukaryotic proteome. The endoplasmic reticulum membrane complex (EMC) plays a critical role in the insertion of membrane proteins into the lipid bilayer of the endoplasmic reticulum (ER) (Chitwood et al., 2018; Guna et al., 2018; Shurtleff et al., 2018). The EMC is ubiquitously expressed and highly conserved (Volkmar and Christianson, 2020; Wideman, 2015). Ten subunits (EMC1-10) have been identified in mammals: EMC1, EMC3-7, and EMC10 are membrane-spanning, whereas EMC2, EMC8, and EMC9 are soluble (EMC8 and EMC9 are structural redundant and mutually exclusive) (**Figure 1A**) (O’Donnell et al., 2020). Recent cryo-electron microscope (cryo-EM) structures showed the overall similar organization of the human EMC (Pleiner et al., 2020) and yeast EMC (Bai et al., 2020): both have a large ER luminal region, 12 transmembrane helices, and a smaller cytosolic region. Research has shown that EMC1-3, 5, and 6 are the core subunits since their depletion leads to co-translational degradation of other subunits, malfunction in the assembly of the full mature EMC, and loss of EMC’s overall activity (Volkmar et al., 2019).

**Figure 1.**
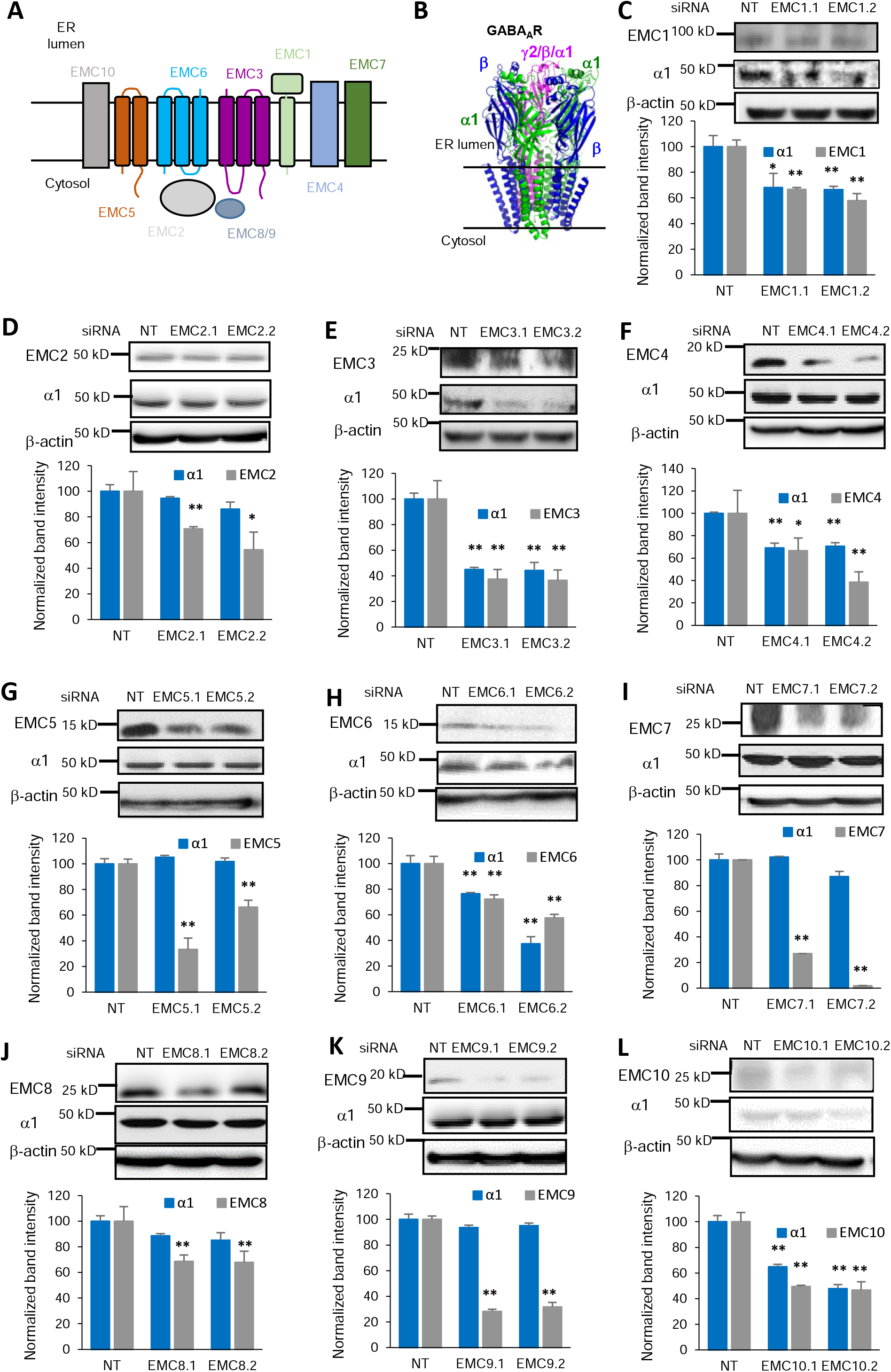
Effect of depleting individual EMC subunits on endogenous GABA_A_ receptor α1 subunit protein levels. **(A)** Schematics of ten EMC subunits. **(B)** Cartoon Representation of heteropentameric GABA_A_ receptors. The most common subtype in the mammalian central nervous system consists of α1, β2/β3, and γ2 subunits. **(C to L)** Endogenous total GABA_A_ receptors α1 subunit’s protein level change upon knocking down individual EMC subunits. Mouse hypothalamus GT1-7 neuronal cells were transfected with siRNA against EMC1 to EMC10, respectively. Two distinct siRNAs targeting each of the ten EMC subunits were used to minimize the potential off-target effects. Forty-eight hours post transfection, proteins were extracted and analyzed by western blotting. β-actin was used as the loading control. Normalized band intensity was shown below the images (n = 3). Each data point is presented as mean ± SEM. *, *p* < 0.05; **, *p* < 0.01. NT: Non-targeting scrambled siRNA.

A growing number of EMC-dependent client membrane proteins have been reported (Volkmar and Christianson, 2020). The EMC was first described in yeast in a genetic screen for protein folding factors (Jonikas et al., 2009). Later, the EMC was identified in the screen of the interactome for the mammalian ER-associated degradation (ERAD) network (Christianson et al., 2011). Recently, the EMC’s important role has been recognized in inserting tail-anchored membrane proteins post-translationally and a subset of multi-pass membrane proteins co-translationally. The EMC acts as an insertase for tail-anchored proteins with moderately hydrophobic transmembrane domains (Guna *et al.*, 2018). Subsequently, the EMC was reported to facilitate the insertion of the first transmembrane domain of certain G-protein coupled receptors (GPCRs), such as β-adrenergic receptors (Chitwood *et al.*, 2018). Furthermore, quantitative proteomics analysis comprehensively identified EMC-dependent membrane protein clients, showing that the EMC enables the biogenesis of membrane proteins with destabilizing features, such as polar and charged residue-containing transmembrane domains (Shurtleff *et al.*, 2018; Tian et al., 2019). The idea that the EMC could act as a more general factor to facilitate the biogenesis of membrane proteins is also intriguing since the EMC regulates the function of a variety of membrane proteins, including cystic fibrosis transmembrane conductance regulator (CFTR) in yeast, acetylcholine receptors in *C. elegans*, and rhodopsins and transient receptor potential (TRP) channels in *Drosophila* (Louie et al., 2012; Richard et al., 2013; Satoh et al., 2015).

We focus on proteostasis maintenance of neurotransmitter-gated ion channels (Fu et al., 2016). Due to the nearly exclusive expression of such membrane proteins in the central nervous system, previous screenings in yeast and mammalian cells did not identify the potential involvement of the EMC in their biogenesis (Shurtleff *et al.*, 2018; Tian *et al.*, 2019). Interestingly, EMC6 was shown to positively regulate the expression of acetylcholine receptors and the response to GABA in *C. elegans* (Richard et al., 2013). Therefore, here we evaluated the role of the EMC in the biogenesis of GABA_A_ receptors, the primary inhibitory ion channels in the mammalian central nervous system (Akerman and Cline, 2007). Functional GABA_A_ receptors are assembled as a pentamer in the ER from eight subunit classes: α1-6, β1-3, γ1-3, δ, ε, θ, π, and ρ1-3 (Sequeira et al., 2019). The most common subtype in the human brain contains two α1 subunits, two β2/β3 subunits, and one γ2 subunit (**Figure 1B**) (Laverty et al., 2019; Zhu et al., 2018). Individual subunits need to fold into their native structures in the ER and assemble with other subunits correctly on the ER membrane to form a heteropentamer (Barnes, 2001; Jacob et al., 2008; Martenson et al., 2017). Only properly assembled receptors exit the ER, traffic through the Golgi for complex glycosylation, and reach the plasma membrane for their function (Lorenz-Guertin and Jacob, 2018). The proteostasis network, which contains chaperones (such as BiP and calnexin), ERAD factors (such as Hrd1, Sel1L, and VCP), and trafficking factors (such as LMAN1), orchestrates the folding, assembly, degradation, and trafficking of GABA_A_ receptors, which is essential for their function (Di et al., 2013; Di et al., 2016; Fu et al., 2019; Wang et al., 2013). Loss of function of GABA_A_ receptors causes a variety of neurological diseases, including epilepsy and intellectual disability (Hernandez and Macdonald, 2019). Furthermore, numerous clinical variants in a single subunit cause subunit protein misfolding and/or disrupt assembly of the pentameric complex in the ER (Fu et al., 2022). Such variant subunits are retained in the ER and subject to excessive ERAD, being dislocated into the cytosol and degraded by the proteasome. This leads to decreased cell surface localization of the receptor complex and imbalanced neural circuits. Therefore, operating the proteostasis network of GABA_A_ receptors has the promise to fine-tune their functional surface expression to ameliorate related neurological diseases (Fu et al., 2018; Wang et al., 2014).

Here we aim to understand the role of each individual subunit of EMC on the proteostasis maintenance of GABA_A_ receptors. We found that EMC3 and EMC6 have the most prominent effect on the protein levels of endogenous GABA_A_ receptors. Furthermore, the interactions between the EMC and GABA_A_ receptors are dependent on the transmembrane domains. Overexpressing the EMC is sufficient to restore the function of disease-associated variants of GABA_A_ receptors.

## RESULTS

### Knocking down EMC3 and EMC6 reduces the protein levels and function of endogenous GABA_A_ receptors

We evaluated the effect of individual EMC subunits on endogenous GABA_A_ receptor protein levels using a mouse GT1-7 hypothalamic neuronal cell line, which is one of the very few neuronal cells that express endogenous, functional GABA_A_ receptors (with α1 and β3 subunits) (Hales et al., 1992). Additionally GT1-7 is a mature mouse hypothalamic gonadotropin releasing hormone cell line that responses to membrane depolarization (Hales et al., 1994), which is a key characteristic of neurotransmitter-gated ion channels. To investigate whether there are key subunits of the EMC that play critical roles in the biogenesis of GABA_A_ receptors, we conducted siRNA screening by using two distinct siRNAs targeting each of the ten EMC subunits to minimize the potential off-target effects. To increase knockdown efficiency, we performed two siRNA transfections at 24 hr and 48 hr before harvesting the proteins for SDS-poly acrylamide gel electrophoresis (PAGE) and Western blot analysis (**Figure 1C to 1L**).

Among the five core subunits of the EMC (EMC1-3, 5, and 6) that are critical for its assembly and activity, EMC1, EMC3 and EMC6 positively regulated α1 protein levels (**Figure 1C**, **1E**, and **1H**), whereas the knockdown of EMC2 or EMC5 did not influence the α1 protein levels significantly (**Figure 1D** and **1G**), suggesting that the formation of the mature EMC is not a prerequisite for the regulation of GABA_A_ receptor protein levels. The most substantial reduction of the total α1 protein levels was observed with the depletion of EMC3 (*p* < *0.01*, **Figure 1E**) and EMC6 (*p* < *0.01*, **Figure 1H**). This is noted as decreased band intensity for EMC3.1 and EMC3.2, compared to the non-targeting (NT) control, to 38% and 37% respectively (**Figure 1E**). Correspondingly, when α1 was detected, two similar decreased band intensities were observed as well, to 41% and 45% respectively (**Figure 1E**). We also noted significant effects on EMC6 knockdown (*p* < 0.01, **Figure 1H**) as well. This is noted as decreased band intensity for EMC6.1 and EMC6.2, compared to NT, to 70% and 58% respectively. Correspondingly, when α1 was detected, two similar decreased band intensities were observed as well, to 75% and 35% respectively **(Figure 1H**). Therefore, our result suggests that individual EMC subunit, such as EMC3 and EMC6, is sufficient to regulate GABA_A_ receptor biogenesis (also see below). EMC3 is the only EMC subunit that exhibits homology to Oxa1 family proteins, which are known membrane protein insertases (Volkmar and Christianson, 2020; Wideman, 2015), and EMC6 plays a critical role in regulating the biogenesis of acetylcholine receptors in *C. elegans* (Richard *et al.*, 2013).

Among the five peripheral subunits of EMC (EMC4, 7-10) that are not essential for the complex’s stability or assembly, EMC4 and EMC10 positively regulated α1 protein levels (**Figure 1F** and **1L**), whereas the knockdown of EMC7-9 did not influence the α1 protein levels significantly (**Figure 1I**, **1J**, and **1K**), again suggesting a subunit-specific contribution of the EMC. It is worth noting that all cytosolic soluble EMC subunits (EMC2, 8-9) did not influence α1 protein levels significantly, suggesting that the transmembrane domains in other membrane-spanning EMC subunits could play a critical role in regulating GABA_A_ receptors biogenesis in the ER membrane (also see below).

Based on GABA_A_ receptors α1 subunit results (**Figure 1)**, we performed additional experiments to detect endogenous β3 protein levels from EMC3 or EMC6 knockdown samples (**Figure 2A**) to further confirm their effects on other subunits of GABA_A_ receptors. This is noted as decreased band intensity for EMC3.1 and EMC3.2, compared to NT, to 50% and 25% respectively. Correspondingly, when β3 was detected, two similar decreased band intensities were observed as well, to 55% and 30% respectively (**Figure 2A**, left panel). We also noted significant effects on EMC6 knockdown (*p* < 0.01 **Figure 2A**) as well. This is noted as decreased band intensity for EMC6.1 and EMC6.2, compared to NT, to 55% and 30% respectively. Correspondingly, when β3 was detected, two similar decreased band intensities were observed as well, to 50% and 25% respectively (**Figure 2A**, right panel).

**Figure 2.**
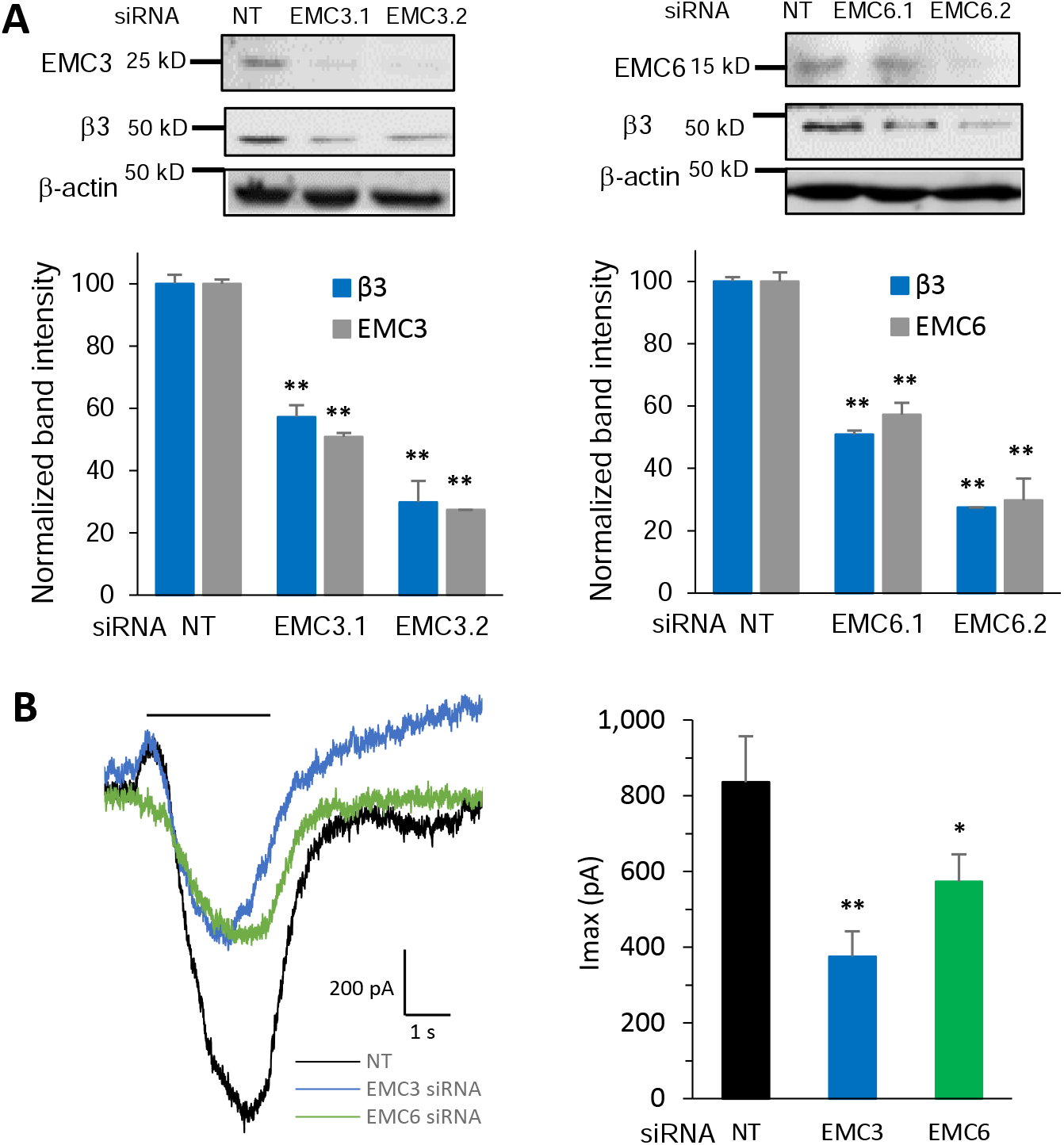
Significant reduction of endogenous GABA_A_ receptor β3 subunits protein levels and whole-cell patch-clamping currents were observed from knocking down EMC3 and EMC6. Mouse GT1-7 neurons were incubated with siRNA of EMC3 or EMC6 for 48 hours. **(A)** Proteins were extracted and analyzed by western blotting; normalized band intensity was shown below the images (n = 3), with β-actin as the loading control. **(B)** Whole-cell patch-clamping was performed on the cells with the IonFlux Mercury 16 ensemble plates at a holding potential of −60 mV. GABA (1 mM) was applied for 4 s, as indicated by the horizontal bar above the currents. The peak currents (Imax) were acquired and analyzed by Fluxion Data Analyzer (n = 8 - 16). Each data point is presented as mean ± SEM. *, *p* < 0.05; **, *p* < 0.01. NT: Non-targeting scrambled siRNA; pA, picoampere.

To understand their effects on the function of endogenous GABA_A_ receptors, we carried out whole-cell patch-clamping electrophysiological recording using GT1-7 neurons after siRNA treatment of EMC3 and EMC6. The peak GABA-induced current in the non-targeting siRNA-treated GT1-7 neurons is 837 pA; consistently, knocking down EMC3 or EMC6 decreased the peak current (Imax) to 375 pA and 573 pA, corresponding to 55% and 32% reduction respectively (**Figure 2B**). These results unambiguously demonstrated EMC3 and EMC6 as key EMC subunits to positively regulate protein expression and function of endogenous GABA_A_ receptors.

### EMC3 and EMC6 promote anterograde trafficking of GABA_A_ receptors

Since GABA_A_ receptors need to traffic to the plasma membrane for their function, we further tested the hypothesis about the necessity of the EMC with respect to surface trafficking of endogenous GABA_A_ receptors using immunofluorescence staining in primary rat cortical neurons **(Figure 3A)**. EMC3 and EMC6 depletion was achieved through the lentivirus transduction of four siRNA pools (Ding and Kilpatrick, 2013). At day-in-vitro (DIV) 6 of the neurons, lentivirus transduction was carried out at a multiplicity-of-infection (MOI) of 30, that is, the ratio of lentivirus count to neuron cells count in each well. Immunofluorescence staining was performed at DIV 12 (Glynn and McAllister, 2006). The application of an anti-GABA_A_ receptor subunit antibody that recognizes the extracellular epitope without a prior membrane permeabilization step using detergent enables us to label the cell surface proteins of α1, β2/3 and γ2 subunits. Application of the Alexa Fluor Dye-conjugated secondary antibody enables the imaging of the surface subunits using a confocal laser-scanning microscope. Clearly, depletion of EMC3 and EMC6 reduced the surface expression of α1 subunits by 37% and 41%, β2/3 subunits by 61% and 50%, and γ2 subunits by 44% and 58% in cortical neurons **(Figure 3A)**, indicating that EMC3 and EMC6 play a critical role in positively regulating the surface trafficking of all major subunits of endogenous GABA_A_ receptors.

**Figure 3.**
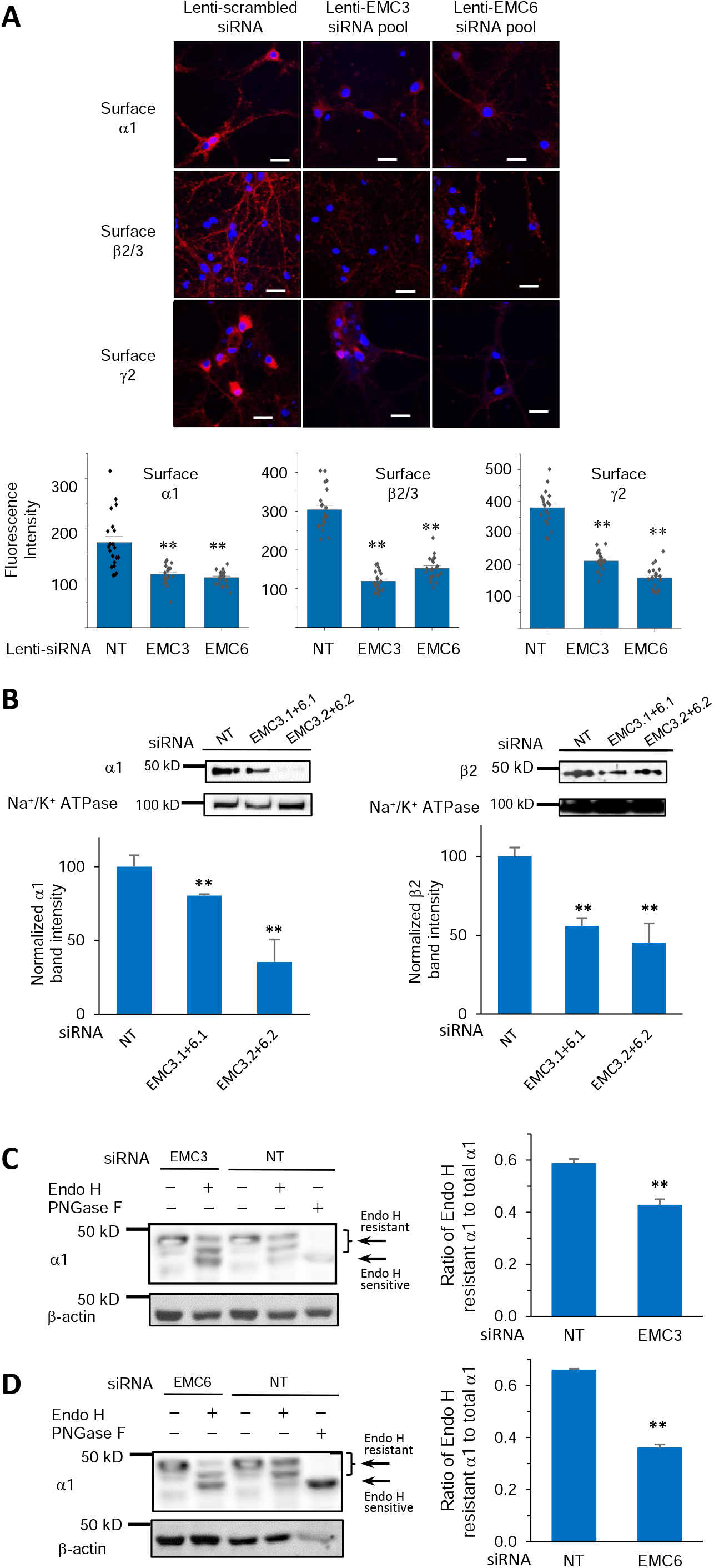
EMC3 and EMC6 promote anterograde trafficking of GABA_A_ receptors. **(A)** Confocal microscopy imaging of primary rat cortical neurons demonstrated reduced surface expression of GABA_A_ receptors after siRNA treatment of EMC3 and EMC6 through lentivirus transduction. Lentivirus were generated from transiently transfected HEK293T cells with the following plasmids and collected after 60 h from the media passing through 0.45 μm filter: EMC3- or EMC6-set of four siRNA lentivectors, packaging and envelop plasmids. Furthermore, the lentivirus were concentrated and quantified with a qPCR lentivirus titration. At day-in-vitro (DIV) 6 of the primary rat cortical neurons, lentivirus transduction was carried out at a multiplicity-of-infection (MOI) of 30. At DIV 12, neurons were stained for cell surface GABA_A_ receptor α1 subunits (top row), β2/β3 subunits (middle row), and γ2 subunits (bottom row), colored in red. DAPI staining for the nucleus was colored in blue. Scale bar = 20 μm. Quantification of the fluorescence intensity by using ImageJ was shown on the bottom after background correction from 20 – 30 neurons. **(B)** Significant reduction of cell surface α1 and β2 subunits of GABA_A_ receptors was observed when both EMC3 and EMC6 were knocked down. We carried out siRNA transfection in HEK293T cells stably expressing α1β2γ2 GABA_A_ receptors. To test the surface expression of GABA_A_ receptors, biotinylation experiments were performed after 48 h post siRNA transfection of both EMC3 and EMC6. Surface proteins were enriched through biotin-neutravidin affinity purification, and Western blot analysis was applied to detect surface α1 and β2 subunits. Na^+^/K^+^ ATPase served as loading control of cell surface proteins. Normalized band intensity was shown below the images (n = 3). **(C, D)** EMC3 and EMC6 promote GABA_A_ receptors’ trafficking from ER to Golgi as demonstrated through endoglycosidase H (Endo H) digestion. EMC3 or EMC6 siRNA transfection was applied in HEK293T cells stably expressing α1β2γ2 GABA_A_ receptors; 48 h post transfection, proteins were extracted, and subjected to Endo H digestion and Western blot analysis. Endo H resistant bands (top two bands in lanes 2 and 4) represent proteins that have correctly folded in the ER, trafficked to Golgi and fully modified with the *N*-linked complex glycans, thus becoming resistant to Endo H. On the other hand, acting upon proteins remaining in ER, Endo H may remove the high mannose structure after the asparaginyl-*N*-acetyl-D-glucosamine on the α1 subunits, generating Endo H sensitive bands (bottom band in lanes 2 and 4). The Peptide-N-Glycosidase F (PNGase F) enzyme treated samples served as a control for unglycosylated α1 subunits (lane 5). Quantification of the ratio of Endo H resistant / total α1 band intensity, as a measure of the trafficking efficiency of α1 subunits, was shown on the right (n = 3). Each data point is presented as mean ± SEM. *, *p* < 0.05; **, *p* < 0.01. NT: Non-targeting scrambled siRNA.

In addition, we carried out surface biotinylation experiments to evaluate the effect of the EMC on exogenously expressed human α1β2γ2 GABA_A_ receptors in HEK293T cells. After 48 h of siRNA transfection of both EMC3 and EMC6, surface proteins were enriched through biotin-neutravidin affinity purification (see Materials and Methods), and Western blot analysis was applied to detect α1 and β2 subunits **(Figure 3B)**. Significant reduction of cell surface α1 subunits of GABA_A_ receptors was observed when both EMC3 and EMC6 were knocked down comparing to NT, to 78% or 34% when using two sets of siRNAs **(Figure 3B, left panel)**. Similarly, cell surface β2 subunits decreased to 55% or 42% when using two sets of siRNAs **(Figure 3B, right panel)**, when both EMC3 and EMC6 were knocked down.

We hypothesized that knocking down EMC3 and EMC6 would lead to folding and assembly defects of GABA_A_ receptors in the ER, and thus their retention in the ER and compromised anterograde trafficking. We demonstrated such effects on GABA_A_ receptors’ trafficking from ER to Golgi through endoglycosidase H (Endo H) digestion. The human α1 subunit of GABA_A_ receptors has two *N*-linked glycosylation sites at N38 and N138. Endo H resistant bands represent glycoproteins that have correctly folded in the ER, trafficked to Golgi and fully modified with the *N*-linked complex glycans, thus becoming resistant to Endo H. On the other hand, acting upon glycoproteins remaining in ER, Endo H may remove the high mannose structure after the asparaginyl-*N*-acetyl-D-glucosamine on the α1 subunits of GABA_A_ receptors. We carried out EMC3 or EMC6 siRNA transfection in HEK293T cells expressing α1β2γ2 GABA_A_ receptors; 48 h post transfection, proteins were extracted, and subjected to Endo H digestion and Western blot analysis **(Figure 3C** and **3D)**. The Peptide-N-Glycosidase F (PNGase F) enzyme treated samples served as a control for unglycosylated α1 subunits (**Figure 3C** and **3D**, lane 5). Two endo H resistant α1 bands were observed, corresponding to singly or doubly glycosylated α1 at N38 and N138 ((**Figure 3C** and **3D**, lanes 2 and 4). Ratio of Endo H resistant α1 to total α1 represents its ER-to-Golgi trafficking efficiency. Such ratio decreased to 0.43 with EMC3 siRNA (**Figure 3C**, lane 2), comparing to a ratio of 0.59 of NT (**Figure 3C**, lane 4). Similarly, the ratio decreased even further to 0.37 with EMC6 siRNA (**Figure 3D**, lane 2), comparing to a ratio of 0.65 of NT (**Figure 3D**, lane 4). Therefore, EMC3 and EMC6 have shown capability of promoting anterograde trafficking of GABA_A_ receptors from the ER to the Golgi (**Figure 3C** and **3D**) and onward to the plasma membrane (**Figure 3A** and **3B**).

### Co-immunoprecipitation from primary cortical neurons demonstrated endogenous interactions between EMC3/EMC6 and a number of major neuroreceptors, including GABA_A_ receptors, N-methyl-D-aspartate receptors (NMDARs), and nicotinic acetylcholine receptors (nAChRs)

To facilitate the biogenesis of its client membrane proteins, the EMC is expected to interact with them. Therefore, we continued to understand the interactions between EMC3/6 and GABA_A_ receptors. Rat cortical neurons at DIV 12 were harvested to perform co-immunoprecipitation. Endogenous interactions were confirmed between α1 subunits and EMC3/EMC6 **(Figure 4A)**. In addition, as expected, pulling down α1 subunits led to the detection of β2/3 and γ2 subunits of GABA_A_ receptors, indicating the endogenous interactions within the pentameric receptors. Further, due to the important role of the proteostasis network in orchestrating GABA_A_ receptor biogenesis, we observed α1-interacting chaperones, including BiP and calnexin, and ERAD factors, including Grp94 and VCP (**Figure 4A**). Previously, we demonstrated their individual roles in assisting the folding and degradation of GABA_A_ receptors (Di *et al.*, 2013; Di *et al.*, 2016; Han et al., 2015a; Han et al., 2015b).

**Figure 4.**
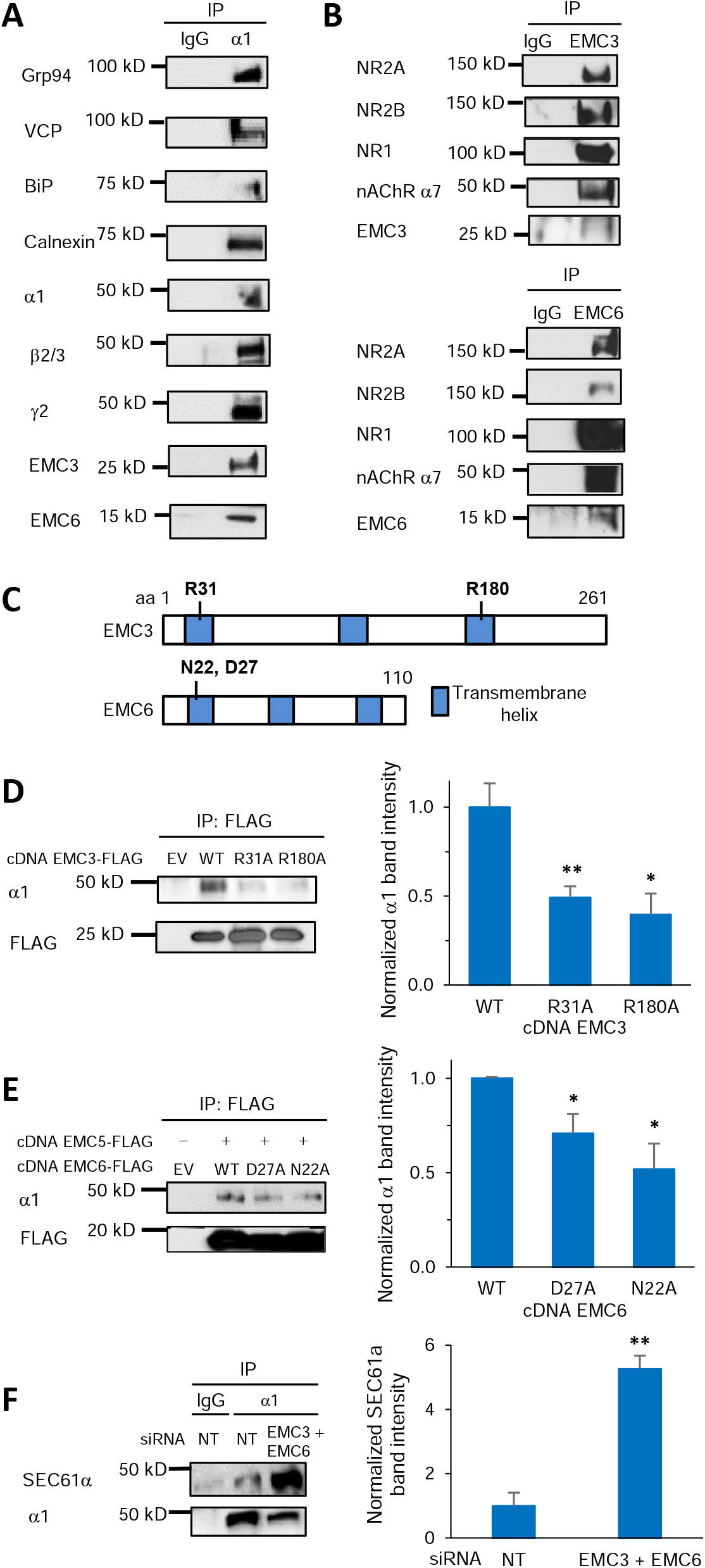
Interactions of EMC3 and EMC6 with neurotransmitter-gated ion channels. **(A)** Co-immunoprecipitation (Co-IP) from primary rat cortical neurons demonstrated endogenous interactions between α1 subunits of GABA_A_ receptors and EMC3, EMC6, and a number of α1-interacting chaperones (BiP and calnexin) and ERAD factors (Grp94 and VCP). Neurons were plated onto 10-cm dishes at a density of 1 million per dish. At DIV 12, proteins were extracted for Co-IP. IgG was used as a negative control during the immunoprecipitation. n = 3. **(B)** Co-IP from primary rat cortical neurons demonstrated endogenous interactions between EMC3/EMC6 and a number of ion channels, including N-methyl-D-aspartate receptors (NMDARs, including NR1, NR2A and NR2B subunits) and nicotinic acetylcholine receptors (nAChR α7 subunit). n = 3. **(C)** Schematic of the primary sequence of EMC3 and EMC6. R31 and R180 in EMC3 and N22 and D27 in EMC6 were reported to influence the biogenesis of EMC-dependent client proteins. **(D)** Mutation of R31A or R180A in EMC3 significantly reduced the interaction of EMC3 with GABA_A_ α1 subunits. The cDNAs of FLAG-tagged EMC3, either in the wild type (WT) form or carrying appropriate mutations of R31A or R180A, were transiently transfected in HEK293T cells stably expressing α1β2γ2 GABA_A_ receptors. 48 h post transfection, proteins were extracted from cell lysates and incubated with anti-FLAG M2 magnetic beads. The immuno-purified eluents were separated through SDS-PAGE gel, and Western blot analysis was performed to detect α1 subunits and FLAG. Quantification of the band intensity of α1 over FLAG post immunoprecipitation was shown on the right (n = 3). **(E)** Mutation of D27A or N22A in EMC6 significantly reduced the interaction of EMC6 with GABA_A_R α1 subunits. Transfection of cDNAs was applied similarly as in **D**, however with co-application of FLAG-tagged EMC5 and EMC6 variants in HEK293T cells stably expressing α1β2γ2 GABA_A_ receptors. Co-IP and visualization of protein bands were carried out the same way as in **D** as well. Quantification of the band intensity of α1 over FLAG-tagged EMC6 post immunoprecipitation was shown on the right (n = 3). **(F)** Significant increase of the interaction of SEC61a and α1 subunits of GABA_A_ receptors was observed when both EMC3 and EMC6 were knocked down. We carried out siRNA transfection in HEK293T cells stably expressing α1β2γ2 GABA_A_ receptors; 48 h post transfection, proteins were extracted from cell lysates and incubated with anti-α1 antibody. The immuno-purified eluents were separated through SDS-PAGE gel, and Western blot analysis was performed to detect SEC61α and α1 subunits. Quantification of the band intensity of SEC61α over α1 post immunoprecipitation was shown on the right (n = 3). Each data point is presented as mean ± SEM. *, *p* < 0.05; **, *p* < 0.01. NT: Non-targeting scrambled siRNA; IP: immunoprecipitation; EV: empty vector; WT: wild type.

Furthermore, we evaluated whether the EMC has a more general role in interacting with other major endogenous neurotransmitter-gated ion channels, including nAChRs and NMDA receptors. Both GABA_A_ receptors and nAChRs belong to the Cys-loop superfamily neuroreceptors, sharing pentameric scaffold (Fu *et al.*, 2016), whereas NMDA receptors are tetramers, consisting of two NR1 subunits and two NR2 subunits (NR2A or NR2B, or both) (Hansen et al., 2021). Dysfunction of these receptors leads to neurological, cognitive and psychiatric brain diseases (Shen et al., 2016a; Steinlein, 2012; Wanamaker et al., 2003). Co-immunoprecipitation experiments demonstrated that pulling down EMC3 or EMC6 resulted in the detection of NR1, NR2A, and NR2B subunits of NMDA receptors as well as α7 subunits of the nAChRs in primary cortical neurons **(Figure 4B)**. These experiments successfully confirmed the interactions between the EMC and a number of major neuroreceptors, suggesting that the EMC not only assists folding of GABA_A_ receptors but also potentially plays critical roles for the biogenesis of other multi-pass-transmembrane receptors in the central nervous system.

### Mutation of key transmembrane residues of EMC3 and EMC6 significantly impairs their capability of interacting with GABA_A_ receptors

Next we determined whether the interactions between EMC3/6 and GABA_A_ receptors are through their transmembrane domains. For EMC3, previous literature has shown the importance of positive residues at Arg 31 (R31) in transmembrane helix 1 (TM1) and Arg 180 (R180) in transmembrane helix 3 (TM3) (**Figure 4C**), and the absence of such positive residues destabilize post- and co-translational insertion of EMC-dependent substrates, such as squalene synthase and opioid kappa receptor 1 (Pleiner *et al.*, 2020). For that reason, we hypothesized that a neutral residue at such positions would likewise destabilize the interaction between EMC3 and GABA_A_ receptors. FLAG-tagged WT EMC3 or EMC3 carrying appropriate mutations of R31A or R180A were transiently transfected in HEK293T cells stably expressing α1β2γ2 GABA_A_ receptors. The immuno-purified eluents were separated through SDS-PAGE, and Western blot analysis was performed to detect α1 subunits and FLAG. As demonstrated by normalized α1 intensity over FLAG, comparing to WT EMC3, significant decrease of the interaction of mutant EMC3 and α1 subunits was observed, to an extent of 0.48 with R31A, or 0.40 with R180A, respectively **(Figure 4D)**. Therefore, the result indicates that a neutral residue replacement at R31 in TM1 and R180 in TM3 impairs EMC3’ capability to interact with GABA_A_ receptors.

Moreover, EMC6 residues in the hydrophilic vestibule are thought to be necessary for insertion of its substrates (**Figure 4C**) (Pleiner *et al.*, 2020) Asp 27 (D27) in TM1 has been shown with such possibility, and is conserved across several species such as *Homo sapiens, M musculus*, and *S. cerevisiae* (Pleiner *et al.*, 2020). Additionally, Asn 22 (N22) has been identified in TM1 of EMC6 within hydrogen bonding distance of the main chain of EMC5, which provides insights in explaining how its TM1 stabilizes in the lipid bilayer. Therefore similarly, FLAG-tagged EMC5 and appropriate EMC6 variants were transfected in HEK293T cells stably expressing α1β2γ2 GABA_A_ receptors. Co-expression of EMC5 with EMC6 was necessary since EMC5 is required to stably insert EMC6’s TM1 (Pleiner *et al.*, 2020). Comparing to WT EMC6, its variants led to significant decrease of its interaction with α1 subunits, to an extent of 0.72 with D27A, or 0.55 with N22A, respectively **(Figure 4E)**. The result indicates that D27 and N22 in TM1 play critical roles for EMC6’ interaction with GABA_A_ receptors. Taken together, we showed that the interactions between EMC3/6 and GABA_A_ receptors are dependent on the key charged/polar residues in the TM domains of the EMC, consistent with the role of the EMC as an insertase for multi-pass transmembrane proteins.

Furthermore, since the SEC61 translocon is known to play a crucial role in the insertion of secretory and membrane polypeptides into ER co-translationally (Skach, 2009) and the EMC coordinates with SEC61 for the insertion of transmembrane proteins (Chitwood *et al.*, 2018; O’Keefe et al., 2021), we investigated the potential orchestration of SEC61 and EMC on the GABA_A_ receptor biogenesis. We carried out siRNA transfection of both EMC3 and EMC6 in HEK293T cells stably expressing α1β2γ2 GABA_A_ receptors. Co-immunoprecipitation experiments showed that significant increase of the interaction of SEC61α and α1 subunits of GABA_A_ receptors was observed when both EMC3 and EMC6 were knocked down, as demonstrated by normalized SEC61α over α1 intensity comparing to NT, to a remarkable extent of 5.3-fold **(Figure 4F)**. Therefore, the result suggests that upon depletion of EMC3 and EMC6, GABA_A_ receptors would be routed to the SEC61 translocon for the insertion into the lipid bilayer.

### Overexpression of EMC3 and EMC5/6 restores functional surface expression of disease associated variants (DAVs) of GABA_A_ receptors

Based on the above demonstrated results, increasing the EMC expression has the promise to enhance forward trafficking and thus surface expression of GABA_A_ receptors, and ultimately their function as neurotransmitter-gated ion channels. To investigate this hypothesis, we evaluated the role of key EMC subunits in epilepsy-associated GABA_A_ receptors variants, which are known to cause subunit misfolding and reduced surface expression, including α1(D219N), α1(G251D), and α1(P260L) (Fu *et al.*, 2022; Han *et al.*, 2015b; Kodera et al., 2016). We carried out cDNA transfection of EMC3 or co-application of EMC5 and EMC6 in HEK293T cells expressing the variants. With EMC3 overexpression, significantly increased surface expression of α1 subunits of GABA_A_ receptors was observed in HEK293T cells stably expressing α1(D219N)β2γ2 **(Figure 5A)**, α1(G251D)β2γ2 **(Figure 5B)** and α1(P260L)β2γ2 **(Figure 5C)**, to 260%, 285%, and 148% respectively. Similarly, with co-application of EMC5 and EMC6, surface α1 subunits increased to 380%, 235%, and 285% respectively in cells expressing α1(D219N)β2γ2 **(Figure 5A)**, α1(G251D)β2γ2 **(Figure 5B)** and α1(P260L)β2γ2 **(Figure 5C)**.

**Figure 5.**
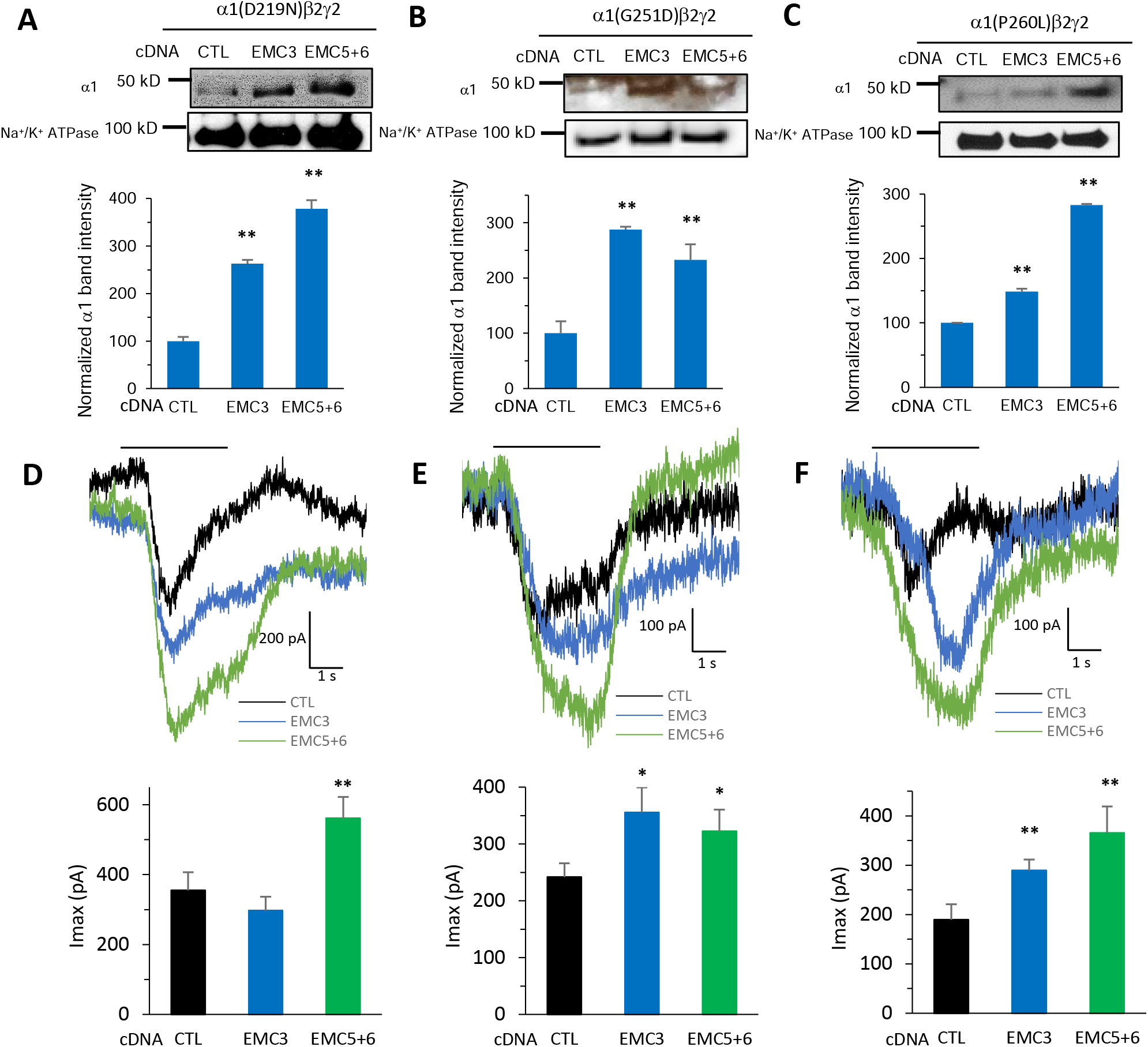
Overexpression of EMC3, and EMC5 and EMC6 restores surface expression and whole-cell currents of disease-associated variants of GABA_A_ receptors. Overexpression of EMC3 and EMC5/6 increased surface expression of α1 subunits of GABA_A_R in HEK293T cells stably expressing α1(D219N)β2γ2 **(A)**, α1(G251D)β2γ2 **(B)** and α1(P260L)β2γ2 **(C).** We carried out cDNA transfection of EMC3, or co-application of EMC5 and EMC6, in corresponding HEK293T cells; 48 h post transfection, surface proteins were enriched through biotin-neutravidin affinity purification, and Western blot analysis was applied to detect α1 subunits. Na^+^/K^+^ ATPase served as loading control of cell surface proteins. Normalized surface α1 band intensity was shown below the images (n = 3). Furthermore, increased whole-cell patch-clamping currents of GABA_A_ receptors were recorded in HEK293T cells stably expressing α1(D219N)β2γ2 **(D)**, α1(G251D)β2γ2 **(E)** and α1(P260L)β2γ2 **(F).** Transfection of cDNA was applied the same way as in **A-C**; 48 h post-transfection, patch clamping was performed on the cells with the IonFlux Mercury 16 ensemble plates at a holding potential of −60 mV. GABA (100 μM) was applied for 4 s, as indicated by the horizontal bar above the currents. The peak currents (Imax) were acquired and analyzed by Fluxion Data Analyzer (n = 10 - 21). Each data point is presented as mean ± SEM. *, *p* < 0.05; **, *p* < 0.01. CTL: Control sample.

To further confirm the EMC’s effects on the function of these variants, whole-cell patch-clamping was performed on cells 48 h post-transfection using the Fluxion automated patch clamping instrument (see Materials and Methods). Increased GABA-induced currents were recorded, as shown in α1(D219N)β2γ2 **(Figure 5D)**, α1(G251D)β2γ2 **(Figure 5E)** and α1(P260L)β2γ2 **(Figure 5F)**. With EMC3 overexpression, the peak current (Imax) increased from 239 pA to 355 pA and from 185 pA to 285 pA respectively in α1(G251D)β2γ2 **(Figure 5E)** and α1(P260L)β2γ2 **(Figure 5F)**; no significant change was observed in α1(D219N)β2γ2 **(Figure 5D)**, potentially due to its lesion being in the ER lumen, while the EMC acts preferably on the transmembrane domains (Chitwood and Hegde, 2019). Moreover, with co-application of EMC5 and EMC6, Imax increased to from 270 pA to 550 pA, 239 pA to 325 pA, and 185 pA to 370 pA respectively in α1(D219N)β2γ2 **(Figure 5D)**, α1(G251D)β2γ2 **(Figure 5E)** and α1(P260L)β2γ2 **(Figure 5F)**. These results highlight the potential of the EMC as a target to enhance the function of DAVs of GABA_A_ receptors.

## DISCUSSION

In this study, we first systemically examined the effects of all 10 individual subunits of the EMC on regulating endogenous GABA_A_ receptor protein expression. Significant reduction of GABA_A_ receptor protein levels and GABA-induced currents were observed from knocking down EMC3 and EMC6 in neurons (**Figures 1–3**). These results suggest a subunit-specific contribution of the EMC in regulating the biogenesis of a multi-pass transmembrane protein, which might not depend on the formation of the complete multi-subunit EMC. Intriguingly, consistent results have been reported that the EMC3-EMC6 fusion protein is sufficient to insert the mitochondrial protein Cox2 and nuclear encoded inner membrane proteins (Güngör et al., 2022), suggesting that certain EMC subunits are capable of carrying out the membrane insertase function. EMC3 is homologous to known membrane protein insertases Oxa1 family proteins (Volkmar and Christianson, 2020; Wideman, 2015), and depletion of EMC3 results in ER stress and activates the unfolded protein response (Huang et al., 2021; Tang et al., 2017). Moreover, EMC3 was reported to coordinate the assembly of lipids and proteins required for surfactant synthesis (Tang *et al.*, 2017), maintain differentiation and function of intestinal exocrine secretory lineages (Huang *et al.*, 2021), and play a critical role in mammalian retinal development (Cao et al., 2021; Zhu et al., 2020). In parallel, in addition to its role in regulating the biogenesis of acetylcholine receptors in *C. elegans* (Richard *et al.*, 2013), EMC6 was reported to regulate the autophagosome formation and knocking down EMC6 impaired autophagy (Li et al., 2013). Moreover, the important role of EMC6 as a therapeutic target for cancer and pancreatic inflammatory diseases has been increasingly recognized (Shen et al., 2016b; Tan et al., 2020; Wang et al., 2017; Xiao et al., 2021).

Our data support a working model about the role of the EMC in GABA_A_ biogenesis in the ER (**Figure 6**). The EMC, including EMC3 and EMC6, interacts with the nascent subunits of GABA_A_ receptors through the transmembrane domains. Mechanistic studies revealed that R31 in TM1 and R180 in TM3 of EMC3, and D27 and N22 in TM1 of EMC6 are essential residues for EMC’s interaction with GABA_A_ receptors (**Figure 4C–4E**), consistent with the structural work of the EMC (Pleiner *et al.*, 2020). Therefore, the EMC facilitates the insertion of the transmembrane domains of GABA_A_ subunits into the lipid bilayer. Subsequently, molecular chaperones, such as BiP and calnexin, promote the productive folding of GABA_A_ receptors (Di *et al.*, 2013; Han *et al.*, 2015b). After the proper assembly of the pentameric receptors on the ER membrane, GABA_A_ receptors engage the trafficking factors, such as LMAN1 (Fu *et al.*, 2019), en route to the Golgi and onward to the plasma membrane. Consistently, we demonstrated that EMC3 and EMC6 promote the anterograde trafficking of GABA_A_ receptors (**Figure 3**). Furthermore, since EMC3 and EMC6 interact with major endogenous neurotransmitter-gated ion channels in primary neurons, including Cys-loop receptors and glutamate receptors **(Figure 4A and 4B)**, the EMC could have a more general role in the central nervous system. It would be of great interest to identify the endogenous interactomes of the EMC, such as EMC3 and EMC6, as a way to determine their client membrane proteins in the central nervous system in the future.

**Figure 6.**
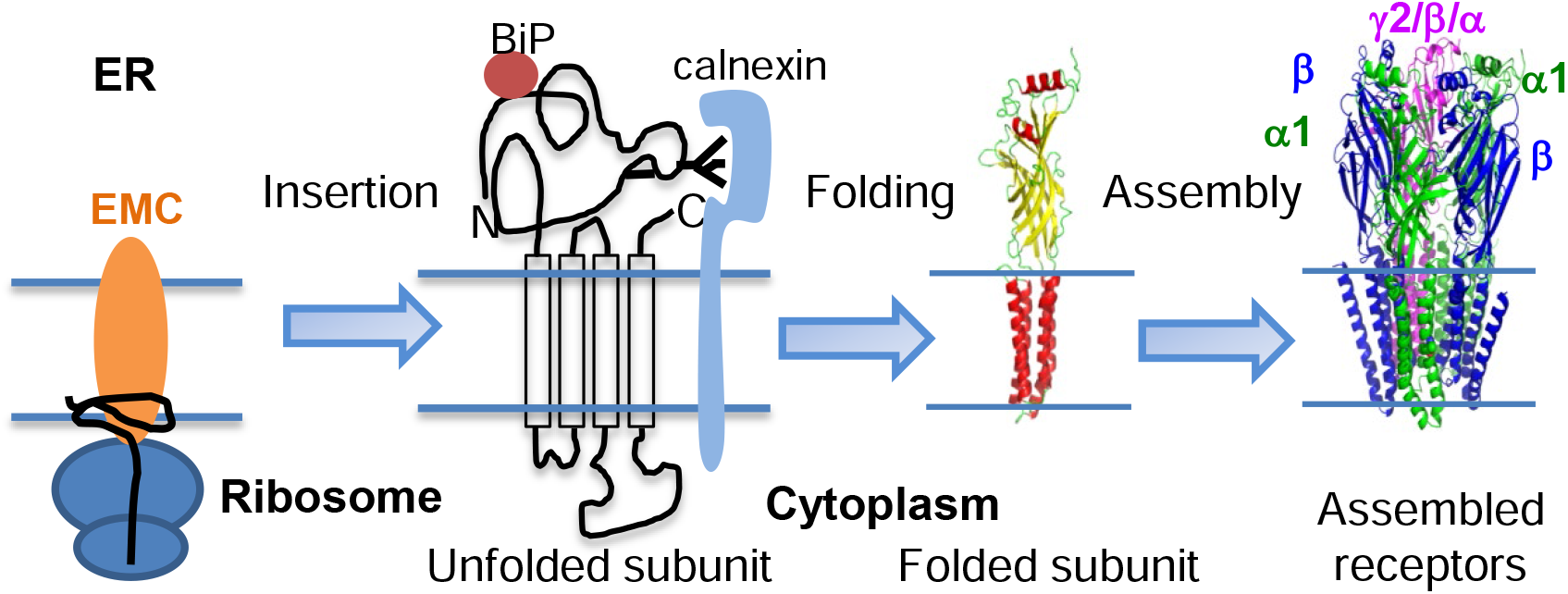
Working model of the role of the EMC on the biogenesis of GABA_A_ receptors in the ER. The EMC facilitates the insertion of the transmembrane domains of GABA_A_ receptor subunits into the lipid bilayer. ER chaperones, such as BiP and calnexin, promote the folding of the ER luminal domains of GABA_A_ receptors for subsequent assembly into pentameric receptors in the ER.

Loss of function of inhibitory GABA_A_ receptors is one of the primary causes of genetic epilepsy. Recent advance in genetics has identified over 150 epilepsy-associated variants in the major subunits (α1, β2, β3, and γ2) of GABA_A_ receptors (Fu *et al.*, 2022; Hernandez and Macdonald, 2019; Hirose, 2014). Despite the development of numerous anti-seizure drugs, about one third epilepsy patients are resistant to current drug treatment (Rincon et al., 2021), and many of them have genetic causes (Smolarz et al., 2021). Therefore, there is an urgent need to develop new therapeutic strategy to treat epilepsy, especially drug-resistant epilepsy. Since one major disease-causing mechanism for loss of function of GABA_A_ receptors is their reduced trafficking to the plasma membrane, one promising approach is to adapt the proteostasis network to restore their surface trafficking and thus function (Di *et al.*, 2013; Di et al., 2021; Fu *et al.*, 2018; Han *et al.*, 2015a; Han *et al.*, 2015b). Here, we demonstrated that overexpression of EMC3 and EMC5/6 enhances the functional surface expression of a number of pathogenic GABA_A_ receptor variants **(Figure 5)**, highlighting the potential of the EMC as a therapeutic target to treat genetic epilepsy resulting from loss of function of GABA_A_ receptors.

## Materials and Methods

### Reagents

The pCMV6 plasmids containing human GABA_A_ receptor α1 (Uniprot no. P14867-1), β2 (isoform 2, Uniprot no. P47870-1), γ2 (isoform 2, Uniprot no. P18507-2) subunits, and pCMV6 Entry Vector plasmid (pCMV6-EV) were obtained from Origene. The human FLAG-tagged EMC3 plasmid was purchased from GenScript (catalog #: OHu03021D). The human FLAG-tagged EMC5 plasmid (catalog #: RC207046) and FLAG-tagged EMC6 plasmid (catalog #: RC215548) were obtained from Origene. The mutations EMC3-R31A, EMC3-R180A, EMC6-D27A, and EMC6-N22A were constructed using QuikChange II site-directed mutagenesis Kit (Agilent Genomics, catalog #: 200523). All cDNA sequences were confirmed by DNA sequencing.

The following small interfering RNA (siRNA) duplexes were obtained from Dharmacon: EMC1.1 (J-059370-09-0005), EMC1.2 (J-059370-10-0005), EMC2.1 (J-049743-09-0005), EMC2.2 (J-049743-10-0005), EMC3.1 (J-056059-09-0005), EMC3.2 (J-056059-11-0005), EMC4.1 (J-046351-09-0005), EMC4.2 (J-046351-10-0005), EMC5.1 (J-041149-09-0005) EMC5.2 (J-041149-11-0005), EMC6.1 (J-047425-10-0005), EMC6.2 (J-047425-12-0005), EMC7.1 (J-051219-09-0005), EMC7.2 (J-051219-11-0005), EMC8.1 (J-046488-09-0005), EMC8.2 (J-046488-10-0005), EMC9.1 (J-046998-09-0005), EMC9.2 (J-046998-10-0005), EMC10.1 (J-041588-11-0005), EMC10.2 (J-041588-12-0005), human EMC3.1 (J-010715-17-0005), human EMC3.2 (J-010715-18-0005), human EMC6.1 (J-014711-19-0005), human EMC6.2 (J-014711-20-0005), and Non-Targeting (NT) siRNA (D-001810-01-20), which was used as a negative control.

### Antibodies

The rabbit polyclonal EMC1 antibody (catalog #: AP10226b), rabbit polyclonal EMC3 antibody (catalog #: AP5782a), rabbit polyclonal EMC4 antibody (catalog #: AP14717a), rabbit polyclonal EMC9 antibody (catalog #: AP5632b), and rabbit polyclonal EMC10 antibody (catalog #: AP5188a) were from Abcepta. The rabbit polyclonal EMC2 antibody (catalog #: 25443-1-AP) and mouse monoclonal EMC8 antibody (catalog #:66547-1-IG) were obtained from Proteintech. The rabbit polyclonal EMC5 antibody (catalog #: PA56905), rabbit polyclonal EMC6 antibody (catalog #: PA-107119), and rabbit polyclonal EMC7 antibody (catalog #: PA5-52688) were from Pierce.

The mouse monoclonal anti-GABA_A_ α1 subunit antibody (clone BD24) (catalog #: MAB339) and mouse monoclonal anti-GABA_A_ β2/β3 antibody (catalog #: 05-474) were obtained from Millipore. The rabbit monoclonal anti-GABA_A_ α1 subunit antibody (catalog #: 224203) and rabbit polyclonal anti-GABA_A_ γ2 antibody (catalog #: 224003) were obtained from Synaptic systems. The rabbit polyclonal anti-GABA_A_ α1 antibody came from R&D systems (catalog #: PPS022). The mouse monoclonal anti-β-actin antibody (catalog #: A1978) and mouse monoclonal FLAG antibody (catalog #: F1804) came from Sigma Aldrich. The fluorescent anti-β-actin antibody Rhodamine came from Biorad (catalog #: 12004163). The rabbit polyclonal anti-calnexin (catalog #: ADI-SPA-860-F) and rat polyclonal anti-Grp94 (catalog #: ADI-SPA-850-F) antibodies were purchased from Enzo Life Sciences. The rabbit polyclonal anti-VCP (catalog #: AP6920b) antibody was obtained from Abgent. The rabbit polyclonal anti-SEC61α antibody was obtained from Proteintech (catalog #: 24935-1-AP). The rabbit polyclonal anti-Grp78 antibody (catalog #: ab21685), rabbit monoclonal anti-NR1 antibody (catalog #: ab109182), rabbit monoclonal anti-NR2A antibody (catalog #: ab124913), rabbit monoclonal anti-NR2B antibody (catalog #: ab183942), and rabbit polyclonal anti-nAChR α7 antibody (catalog #: ab182442), and sodium potassium ATPase antibody (catalog #: ab76020) were purchased from Abcam.

The secondary antibodies used from Invitrogen included: HRP-conjugated goat-anti-rabbit (catalog #: 31460), goat-anti-mouse (catalog #: 31430), and goat-anti-rat (catalog #: 31470), and Alexa Fluor 594-conjugated goat-anti-rabbit (catalog #: A11032) and goat-anti-mouse (catalog #: A11037).

### Cell Culture and transfection

HEK293T cells (catalog #: CRL-3216) were obtained from ATCC, and GT1-7 cells (catalog #: SCC116) were obtained from Millipore. Cells were maintained in Dulbecco’s Modified Eagle Medium (DMEM) (Fisher, catalog #: SH3024301) with 10% heat-inactivated fetal bovine serum (ThermoFisher, catalog #: SH3039603HI) and 1% Penicillin-Streptomycin (Hyclone, catalog #: sv30010) at 37°C in 5% CO_2_. Cells were grown in 6-well plates or 10-cm dishes and allowed to reach ~70% confluency before transient transfection using TransIT-2020 (Mirus, catalog #: 5400), or siRNA treatment (50 nM) using the HiPerfect Transfection Reagent (Qiagen, catalog #: 1029975) according to the manufacturer’s instruction. A second siRNA transfection was performed 24 hours after the first siRNA treatment to increase knockdown efficiency. Forty-eight hours post-transfection, cells were harvested for further analysis.

Stable cell lines for α1β2γ2, α1(D219N)β2γ2, α1(G251D)β2γ2, and α1(P260L)β2γ2 were generated using the G-418 selection method. Briefly, cells were transfected with α1:β2:γ2 (1:1:1), α1(D219N):β2:γ2 (1:1:1), α1(G251D):β2:γ2 (1:1:1) or α1(P260L):β2:γ2 (1:1:1) plasmids, selected in DMEM supplemented with 0.8 mg/mL G418 (Enzo Life Sciences) for 10 days, and then maintained in DMEM supplemented with 0.4 mg/mL G418. G-418 resistant cells were used for experiments.

### Western blot analysis

To harvest total proteins, cells were washed with Dulbecco’s phosphate-buffered saline (DPBS) (Fisher, catalog #: SH3002803). Trypsin (Fisher, catalog #: SH30236.01) was added to lift the cells, and DMEM was added to harvest cells from the dish and pipette into a centrifuge tube. Cells were spun down for 3 minutes at 1000 rpm and then DMEM was removed while avoiding the pellet. Cell pellets were washed with DPBS and centrifuged again at 1000 rpm for 3 minutes. The DPBS was removed, and pellets were stored on ice during transport to a −80°C freezer. Cells were lysed with lysis buffer (50 mM Tris, pH 7.5, 150 mM NaCl, and 2 mM n-Dodecyl-B-D-maltoside (DDM) (GoldBio, catalog #: DDM-5)) supplemented with protease inhibitor cocktail (Roche). Cells were vortexed for 30 seconds followed by ultrasonication for 30 seconds for three times. Then they were centrifuged at 15,000 × *g*, 4 °C for 10 minutes to obtain the supernatant as total proteins. The protein concentration was measured according to Thermo Fisher MicroBCA kit protocol. Cell lysates were loaded with Laemmli sample buffer (Biorad, catalog #: 1610747) with β-mercaptoethanol (1:10 v/v) and separated through SDS-PAGE gel electrophoresis.

Before proceeding to western blot, gels were made ranging from 8% to 20% resolving gel depending on the size of the protein with 4% stacking gel on top. To run SDS-PAGE gel electrophoresis, gels were place in a holding voltage cassette and submerged in the running buffer, which contained the following: 25 mM Tris (Sigma, catalog #: T1503), 192 mM Glycine (Sigma, catalog: # 8898), and 0.1% (w/v) of sodium dodecyl sulfate (SDS, Biorad, catalog #: 1610302). Protein ladder was added (Biorad, catalog #: 1610395). Gels were run at 10 minutes at 100 volts until the samples passed the stacking gel and were uniformly aligned. For the remaining time of 45 min to 1-hour, gels were ran at 150 volts. After running samples to sufficient molecular weight, gels were transferred at 100 volts for 1 hour to a nitrocellulose membrane. The transfer buffer contained the following: 25 mM Tris (Sigma, catalog #: T1503), 192 mM Glycine (Sigma, catalog: # 8898), 20% (v/v) of methanol (Fisher Chemical, catalog #: A452-4). After the transfer, membranes were washed briefly in TBS-T, which contained the following: 20 mM Tris (Sigma, catalog #: T1503), 150 mM NaCl (ACSF, catalog #: S7653), pH 7.6, and 0.1% (v/v) Tween-20 (Sigma, catalog #: P7949-500mL). They were further incubated in 5% non-fat milk powder (Nestle Carnation, catalog #: 43875) in TBS-T for 30 minutes to 2 hours. Following the blocking step, the membranes were incubated in 1% milk with the primary antibody added starting at 1:1000 dilution and adjusted accordingly on subsequent runs. The following day, the membrane was washed 3 times with TBS-T for 10 minutes each and incubated with their secondary antibody (1:10,000) for 1 hour. This was followed by 3 more washed with TBS-T. Afterwards the membrane was exposed with Pico PLUS (catalog #: 34578) or Femto (catalog #: 34096) SuperSignal West chemiluminescent substrates from Thermo Scientific for three minutes. After using different exposure times to get optimal images, results were analyzed to quantify band intensity using ImageJ software from the NIH.

### Co-immunoprecipitation (Co-IP)

Cell lysates (500 μg) were pre-cleared with 30 μl of protein A/G plus-agarose beads (Santa Cruz, catalog #: sc-2003) and 1.0 μg of normal mouse IgG antibody (Santa Cruz, catalog #: sc-2025) for 1 hour at 4°C to remove nonspecific binding proteins. The pre-cleared cell lysates were incubated with 2.0 μg of mouse anti-α1 antibody for 1 hour at 4°C, and then with 30 μl of protein A/G plus agarose beads overnight at 4°C. For FLAG-tagged proteins, the pre-cleared cell lysates were incubated with 30 μl of anti-FLAG M2 magnetic beads (Sigma, catalog #: M8823-5 mL) overnight at 4°C. IgG serves as negative control. The beads were collected by centrifugation at 8000 × g for 30 s or using a magnet separator (Promega), and washed three times with lysis buffer. The complex was eluted by incubation with 30 μl of Laemmli sample loading buffer in the presence of β-mercaptoethanol. The immuno-purified eluents were separated through SDS-PAGE gel, and Western blot analysis was performed.

### Lentivirus transduction in rat cortical neurons

Lentivirus were generated from transiently transfected HEK293T cells and collected after 60 hours from the media. Briefly, HEK293T cells were grown in 10-cm dishes and allowed to reach ~70% confluency before transient transfection using TransIT-2020 (Mirus), according to the manufacturer’s instruction. The following plasmid (6 μg) was added to 10-cm dishes: EMC3-set of four siRNA lentivectors (rat, Abmgood, catalog #: 468690960395), or EMC6-set of four siRNA lentivectors (rat, Abmgood, catalog #: 471140960395), or scrambled siRNA lentivector (Abmgood, catalog #: LV015-G) as the control. Additionally, to form the lentivirus, the following packaging and envelop plasmids were added to all of the 10-cm dishes as well: psPAX2 (6 μg) and pMD2.G (0.75 μg, Addgene, catalog #: 12259). psPAX2 (Addgene plasmid # 12260; http://n2t.net/addgene:12260; RRID:Addgene_12260) and pMD2.G (Addgene plasmid # 12259; http://n2t.net/addgene:12259; RRID:Addgene_12259) were a gift from Didier Trono. Media was changed after 8 h; after 52 additional h, media was harvested and passed through 0.45 μm filter (Advantec, catalog #: 25CS045AS) to collect the lentivirus. Furthermore, the lentivirus were concentrated using Lenti-X concentrator (Takara Bio, Catalog #: 631231) and quantified with the qPCR lentivirus titration kit (Abmgood, catalog #: LV900) according to the manufacturer’s instruction, and saved to −80°C for neuron cells transduction.

Sprague Dawley rat E18 brain cortex tissues were obtained from BrainBits, with provided Hibernate EB (HEB) and NbActiv1 media (Springfield, IL). Prior to plating cells, 10-cm dishes or coverslips (Fisher, catalog #: 12-545-80 CIR-1) in a 24-well plate were coated with 4 mL per plate or 500 μL per well of 50 μg/ml poly-D-lysine (PDL, Sigma, catalog #: P6407) at 4°C overnight. The next day PDL was removed and the plates were washed with sterile distilled water two times. Then they were coated with the same volumes of 5 μg/mL Laminin (Sigma, catalog #: L2020) overnight at 4°C. The plates were then put into a 37°C cell incubator while preparing the neurons for 1 h and Laminin was removed immediately before plating neurons.

Neurons were extracted according to instructions from BrainBits. Briefly, connective tissues of cortex were digested with 2 mL of 2 mg/mL of papain (Brain Bits, catalog #: PAP/HE-Ca) at 30°C water bath for 15 minutes, gently swirling every 5 minutes. Papain was then removed without disturbing the tissue at the bottom of the tube. The HEB media was then added to the tissue tube. The salinized Pasteur pipette was then used to triturate 30 times very slowly and carefully to not add air bubbles with the tip in the middle of the tube. The undispersed pieces were allowed to settle for one minute. The supernatant with dispersed cells were then transferred to a sterile 15 mL tube. This was spun down at 200 × g for one minute and the supernatant was discarded. Afterwards 1-2 mL of NbActiv1 was added while carefully avoiding air bubbles. Neurons were plated at 1 million per 10-cm dish or 25,000 per well of 24-well plate. The neuronal culture media contained the following: 100 mL Neurobasal A (Gibco, catalog #: 21103049), 2 mL B27 (Gibco, catalog #: 17504044), 0.25 mL GlutaMAX (Gibco, catalog #: 35050061) and 1 mL Penicillin-Streptomycin (Hyclone, catalog #: sv30010). Additionally, Ara C (Sigma, catalog #: C6645) was added from day-in-vitro (DIV) 3 at a final concentration of 2 μM each time as half of the media was changed every three days. At DIV 6, lentivirus transduction was carried out to the neurons as indicated; the multiplicity-of-infection (MOI) was at 30, that is, the ratio of lentivirus count to neuron cells count in each well. At DIV 12, neurons were subjected to protein extraction for Co-IP or immunofluorescence staining for confocal microscopy.

### Confocal Immunofluorescence

To label cell surface GABA_A_ receptors, primary neurons that were cultured on coverslips, were fixed with 2% paraformaldehyde in DPBS, blocked with goat serum for 0.5 h at room temperature, and labeled with 100 μL of appropriate anti-α1 (Synaptic Systems, catalog #: 224203), β2/3 (Millipore, catalog #: 05-474), or γ2 (Synaptic Systems, catalog #: 224003) antibodies (1:200) for 1 h without detergent permeabilization. Afterwards, they were incubated at room temperature with 500 μL (1:400) of Alexa 594-conjugated goat anti-rabbit antibody (ThermoFisher, catalog #: A11034), or Alexa 594-conjugated goat anti-mouse antibody (ThermoFisher, catalog #: A11037) for 1h. Finally, cells were permeabilized with saponin (0.2%) for 5 min and incubated with DAPI (1 μg/mL) for 3 min to stain the nucleus. The coverslips were then mounted and sealed. For confocal immunofluorescence microscopy, an Olympus IX-81 Fluoview FV1000 confocal laser scanning system was used. A 60× objective was used to collect images using FV10-ASW software. Quantification of the fluorescence intensity was achieved using the ImageJ software from the NIH.

### Biotinylation of cell surface proteins

To perform the biotinylation assay, 6-cm dishes were coated in a 300-fold dilution of poly-L-lysine (Fisher, catalog #: ICN15017710) at 2 mL per plate for one hour at 37°C. Plates were rinsed twice with DPBS and allowed to air dry. The cells were plated on the coated dishes and incubated for 48 hours post-transfection. The intact cells were washed one time with ice cold DPBS and incubated with the membrane impermeable biotinylation reagent Sulfo-NHS SS-Biotin (0.5 mg/mL; APExBio, catalog #: A8005) in PBS containing 0.1 mM CaCl_2_ and 1 mM MgCl_2_ (PBS+CM) for 30 min at 4 °C to label surface membrane proteins. In order to quench the reaction, the cells were incubated with 1.5 mL of 50 mM glycine in ice cold DPBS-CM twice for 5 minutes at 4°C. They were then washed twice with DPBS. The sulfhydryl groups were then blocked by incubating the cells with 5 nM N-ethylmaleimide (NEM) in DPBS-CM for 15 minutes at room temperature and then the liquid was removed. Cells were scraped off plates and solubilized overnight at 4 °C in lysis buffer (Tris–HCl, 50 mM; NaCl, 150 mM pH 7.5, 2 mM DDM) supplemented with Roche complete protease inhibitor cocktail and 5 mM NEM. The next day the lysates were centrifuged at 16,000 × g for 10 minutes at 4°C to pellet cellular debris. The supernatant was saved, and the protein concentration was measured with a MicroBCA assay. Biotinylated surface proteins were affinity-purified from the above supernatant by incubating for 2 h at 4 °C with 50 μL of immobilized neutravidin-conjugated agarose bead slurry (Fisher, catalog #: PI29200). The samples were then subjected to centrifugation (5,000 × g, 3 min). The beads were washed three times with buffer (Triton X-100, 1%; Tris–HCl, 50 mM; NaCl, 150 mM, pH 7.5) and three times further without Triton X-100. Surface proteins were eluted from beads by incubating for 30 min at room temperature with 80 μL of LSB / Urea buffer (2x Laemmli sample buffer (LSB) with 100 mM DTT and 6 M Urea, pH 6.8) for SDS-PAGE and Western blotting analysis.

### Endoglycosidase H (Endo H) enzyme digestion assay

To remove asparaginyl-*N*-acetyl-D-glucosamine in the N-linked glycans incorporated on the α1 subunit in the ER, total cell lysates were digested with Endo H enzyme (NEBiolab, catalog #: P0703L) with G5 reaction buffer at 37°C for 1h. The Peptide-N-Glycosidase F (PNGase F) enzyme (NEBiolab, catalog #: P0704L) treated samples served as a control for unglycosylated α1 subunits. Treated samples were then subjected to Western blot analysis.

### Automated patch-clamping with IonFlux Mercury 16 instrument

Whole-cell currents were recorded 48 hours post transfection of GT1-7 or HEK293T cells. Automated patch clamping was performed on the Ionflux Mercury 16 instrument (Fluxion Biosciences, California). The extracellular solution (ECS) contained the following: 142 mM NaCl, 8 mM KCl (Sigma, catalog #: P9541), 6 mM MgCl_2_ (Sigma, catalog #: M0250), 1 mM CaCl_2_ (Sigma, catalog #: C3306), 10 mM glucose (Sigma, catalog #: G8270), 10 mM HEPES (Sigma, catalog #: H3375). The intracellular solution (ICS) contained the following: 153 mM KCl (Sigma, catalog #: P9541), 1 mM MgCl_2_ (Sigma, catalog #: M0250), 5 mM EGTA (Sigma, catalog #: E3889), 10 mM HEPES (Sigma, catalog #: H3375). Briefly, cells were grown to 50 to 70 percent confluence on 10-cm dishes. Then 3 mL Accutase (Sigma Aldrich, catalog #: A6964-500mL) was added and the cells were incubated for 3 minutes at 37°C until the cells were floating as observed under microscope with minimal clumps. We then pelleted cells with centrifugation for 1 minute at 200 × g, removed supernatant and resuspended cells in serum free medium HEK293-SFMII (Gibco, catalog #: 11686-029), supplemented with 25 mM HEPES (Gibco, catalog #: 15630-080) and 1% penicillin streptomycin (Hyclone, catalog #: sv30010). Cells were placed on gentle shaking at room temperature for 0.5 to 1 h. Mercury 16 plates were prepared according to manufacture suggestions. Whole-cell GABA-induced currents were recorded at a holding potential of −60 mV, at 100 μM or 1 mM GABA concentration as indicated. The signals were acquired and analyzed by Fluxion Data Analyzer.

### Statistical analysis

All data are presented as mean ± SEM. Statistical significance was evaluated using Student’s *t*-test if two groups were compared and one-way ANOVA followed by post-hoc Tukey test if more than two groups were compared. A *p* < 0.05 was considered statistically significant. *, *p* < 0.05; **, *p* < 0.01.

## Acknowledgements

This work was supported by the National Institutes of Health (R01NS105789 and R01NS117176 to TM, R01NS117176-02S1 NIH Research Diversity Supplement to AW) and Flora Stone Mather Center for Women Research and Professional Development Grant (to AW).

## Author contributions

Conceptualization, AW, YW and TM; Data curation: AW and YW; Formal analysis: AW, YW, and TM; Funding acquisition: AW and TM; Supervision: YW and TM; Writing – original draft: AW, YW, and TM; Writing – review & editing: AW, YW and TM.

